# Patient cerebral organoids capture Alzheimer’s disease proteomic biomarkers and drug targets

**DOI:** 10.64898/2026.06.22.733874

**Authors:** Shannon Thomson, Xinran C. Li, Sophie Walker, Tiffany C.Y. Tang, Mark E. Graham, Sam W.Z. Olechnowicz, Rory Bowden, Global Neurodegeneration Proteomics Consortium, Katherine A. Waugh, Chad Slawson, Jeffrey M. Burns, Russell H. Swerdlow, Heather M. Wilkins, Artur Shvetcov, Caitlin A. Finney

**Author notes:** These authors share senior authorship (Artur Shvetcov, Caitlin A. Finney). Corresponding author. E; A: 176 Hawkesbury Road, Westmead, NSW, Australia 2145.

## Abstract

Patient iPSC-derived cerebral organoids are a leading human model of Alzheimer’s disease, yet their proteome has never been benchmarked against human disease. Clinical cohorts now nominate thousands of biomarkers and drug targets across three proteomic platforms, and whether patient organoids capture these candidates is unknown. Here, we profile AD and control cerebral organoids containing neurons, astrocytes, and microglia on the three platforms driving clinical discovery, mass spectrometry, SomaScan, and Olink, in both conditioned media and lysate. Benchmarked against 121 studies and clinical cohorts of over 17,000 plasma, CSF, and cortex samples, patient organoids detect almost every nominated candidate and reproduce the disease-associated change in roughly one in four of the most reproducible. This convergence spans plasma, CSF, and cortex, and extends to synaptic, mitochondrial, and proteostatic biology. We provide the first multi-platform reference proteome of a patient-derived AD model, establishing it as a translationally relevant system for studying AD.

## INTRODUCTION

Patient induced pluripotent stem cell (iPSC)-derived cerebral organoids have become a leading human model for studying Alzheimer’s disease (AD) ^1–3^. They recapitulate amyloid-β and tau accumulation ^4–8^ and are increasingly used to dissect disease mechanisms and test candidate biomarkers and therapeutics ^1–3^. This has been further supported by the FDA Modernization Act 2.0, which removed the requirement for animal testing in drug development and named organoids among the human models that can provide evidence of drug efficacy and safety ^9–11^. Despite this growing reliance, cerebral organoids models of AD are largely validated on whether they accumulate amyloid-β and tau ^12^. AD, however, is marked by widespread proteomic changes that are central to neurodegeneration and contribute to cognitive decline ^13–15^. Further, recent large-scale clinical cohorts have nominated hundreds to thousands of AD protein biomarkers and drug targets, drawing on harmonized proteomic resources such as the Accelerating Medicines Partnership-AD (AMP-AD) ^16,17^, Alzheimer’s Disease Neuroimaging Initiative (ADNI) ^18,19^, UK Biobank ^20^, and Global Neurodegeneration Proteomics Consortium (GNPC) ^21^. These cohorts largely generate their data on three proteomic platforms, mass spectrometry, SomaScan, and Olink Proximity Extension Assay (PEA) ^17,20,21^, platforms which share only around 9% of cerebrospinal fluid (CSF) and 4% of plasma proteins ^22^ and often disagree even where they overlap ^22–25^. A candidate nominated on one platform therefore cannot be confirmed in a model profiled on another, yet no AD organoid has been profiled on all three. Advancing the translation of these candidates requires a pre-clinical disease model that expresses the relevant protein, reproduces its change in disease, and can be interrogated on the same platforms that nominated it. Whether patient cerebral organoids meet this requirement and can therefore be used to study the biomarkers and drug targets emerging from these cohorts has not been tested.

Here, we benchmark the proteome of patient cerebral organoids against human AD across all three proteomic platforms. We generated AD and control iPSC-derived cerebral organoids containing neurons, astrocytes, and microglia, and profiled both the conditioned media and organoid lysate using label-free mass spectrometry (LC/MS), SomaScan 11k assay, and Olink Explore HT. We then benchmarked the organoid proteome against AD biomarkers and drug targets nominated by 121 published proteomic studies of plasma, CSF, and frontal cortex, and against differentially abundant proteins from clinical cohorts comprising 13,623 plasma, 2,110 CSF, and 1,275 cortex samples. We show that patient organoids detect the majority of clinically nominated AD biomarkers and drug targets and reproduce the disease-associated change in approximately one in four of the most reproducible candidates. This convergence holds across plasma, CSF, and cortex, and extends to the synaptic, mitochondrial, and proteostatic processes that characterize AD. Together, this work establishes patient cerebral organoids as a translationally relevant human model of AD, and provides the field with a multi-platform reference proteome for prioritizing clinically nominated biomarkers and drug targets for mechanistic and therapeutic study.

## RESULTS

### Patient cerebral organoids carry a reproducible AD proteomic signature across three platforms

To map the proteomic landscape of AD in a patient-derived model, we generated cerebral organoids containing neurons, astrocytes, and microglia from AD and control iPSCs. We profiled both conditioned media and organoid lysate using LC/MS, SomaScan 11k assay, and Olink Explore HT (Fig. 1a). We then compared AD biomarkers and drug targets identified in the literature and in large clinical cohorts to those recovered in conditioned media and lysate, respectively (Fig. 1a). This design pairs the organoid secretome with patient biofluids and the lysate with cortical tissue, allowing us to benchmark patient cerebral organoids against each of the proteomic platforms that now drive AD biomarker and drug target discovery.

**Figure 1.**
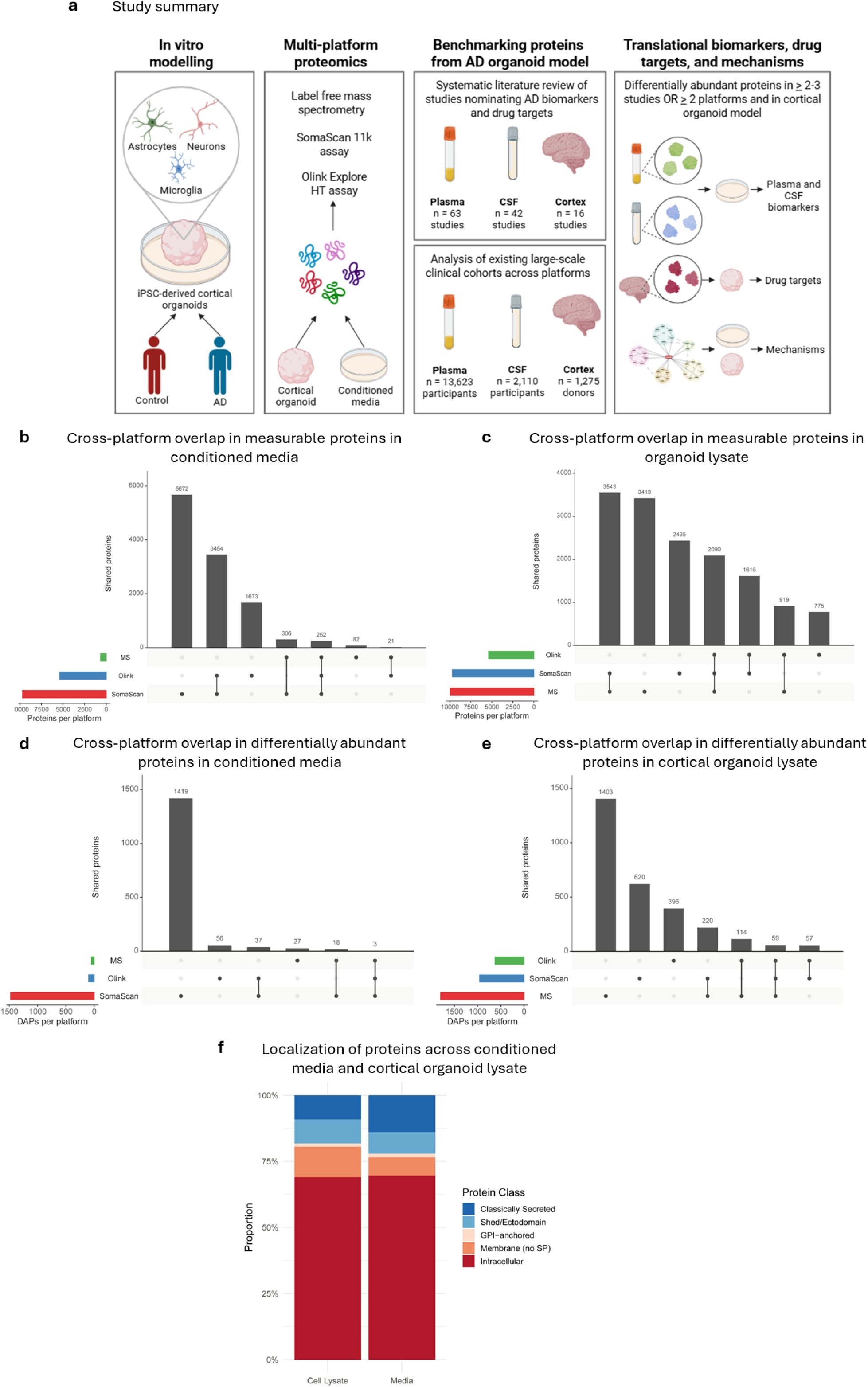
Patient cerebral organoids carry an AD proteomic signature across three proteomic platforms. (**a**) Study design. AD and control iPSC-derived cerebral organoids containing neurons, astrocytes, and microglia were generated, and both the conditioned media and organoid lysate were profiled using label-free mass spectrometry (LC/MS), the SomaScan 11k assay, and Olink Explore HT. AD biomarkers and drug targets nominated in the published literature and in large clinical cohorts were then benchmarked against those recovered in the conditioned media and lysate, respectively. (**b**) UpSet plot of proteome coverage across LC/MS, SomaScan, and Olink in the conditioned media, with 252 proteins shared across all three platforms. (**c**) UpSet plot of proteome coverage across LC/MS, SomaScan, and Olink in the organoid lysate, with 2,090 proteins shared across all three platforms. (**d**) Differentially abundant proteins (DAPs; FDR < 0.05) between AD and control organoids in the conditioned media, collapsed into three tiers by the number of platforms supporting each call: Tier 1 (all three platforms; 3 DAPs), Tier 2 (two platforms; 55 DAPs), and Tier 3 (a single platform; 1,502 DAPs). (**e**) DAPs (FDR < 0.05) between AD and control organoids in the organoid lysate, collapsed into the same three tiers: Tier 1 (59 DAPs), Tier 2 (391 DAPs), and Tier 3 (2,419 DAPs). (**f**) Predicted subcellular localization of DAPs detected by any platform in the conditioned media and organoid lysate. Intracellular proteins accounted for approximately 70% of DAPs in both sample types, with the overall class distribution differing between media and lysate (Fisher’s exact test, *p* = 5×10^-4^). Relative to lysate, media DAPs were enriched for classically secreted proteins (14.0% versus 9.1%) and depleted of membrane proteins lacking a signal peptide (6.9% versus 11.6%). Abbreviations: AD: Alzheimer’s disease; DAP: differentially abundant protein; iPSC: induced pluripotent stem cell; LC/MS: liquid chromatography mass spectrometry.

We first characterized the proteome coverage of each platform across both sample types. In conditioned media, LC/MS detected 661 proteins, SomaScan detected 9,684 proteins after removal of proteoforms and uncharacterized targets from 11,049 features, and Olink detected 5,399 proteins (Supplementary Table 1). Only 252 media proteins were shared across all three platforms, with the largest pairwise overlap between the two affinity platforms (3,454 proteins shared between SomaScan and Olink, compared with 306 between LC/MS and SomaScan and 21 between LC/MS and Olink) (Fig. 1b). In the organoid lysate, LC/MS coverage expanded markedly to 9,971 proteins, approaching the depth of the affinity platforms. Cross-platform coverage similarly rose, with 2,090 proteins shared across all three platforms (Fig. 1c). Here, the largest pairwise overlap shifted to LC/MS and SomaScan (3,543 proteins), followed by Olink and SomaScan (1,616) and Olink and LC/MS (919) (Fig. 1c). This limited overlap reflects the differing coverage of the platforms rather than a failure of any one of them, as each measures a largely distinct set of proteins, fixed by panel content for the affinity assays and by detectability for LC/MS ^23–27^. This mirrors the coverage differences seen across clinical cohorts and underscores why a model must be profiled on all three platforms to be benchmarked against each of them ^22^.

To identify AD-associated changes, we examined differentially abundant proteins (DAPs) between AD and control organoids. Across all six platform and sample type combinations, DAPs separated samples by disease status on principal component analysis (PCA), indicating a reproducible AD signal at the proteome level (Supplementary Fig. 1a-f). At FDR < 0.05, the conditioned media yielded 48 DAPs by LC/MS, 1,546 by SomaScan, and 96 by Olink (Supplementary Fig. 2a-c). The organoid lysate yielded substantially more, with 1,796 by LC/MS, 1,007 by SomaScan, and 626 by Olink (Supplementary Table 1; Supplementary Fig. 2d-f; Fig. 1d, e). To isolate the most reproducible changes, we collapsed DAPs into three tiers defined by the number of platforms supporting each call. In the conditioned media, three DAPs were supported by all three platforms (Tier 1), namely tenascin C (TNC), prostaglandin D2 synthase (PTGDS), and growth differentiation factor 15 (GDF15), while 55 were supported by two platforms (Tier 2) and 1,502 by a single platform (Tier 3) (Fig. 1d). The organoid lysate followed the same structure, with 59 Tier 1, 391 Tier 2, and 2,419 Tier 3 DAPs (Fig. 1e). The small number of Tier 1 DAPs follows directly from the limited cross-platform coverage, as a protein can reach Tier 1 only if it is both measured and differentially abundant on all three platforms.

We next asked whether the two sample types captured biologically distinct fractions of the AD proteome by annotating each DAP detected by any platform according to its predicted subcellular localization. Intracellular proteins dominated both sample types at approximately 70%, consistent with substantial intracellular protein turnover into the media (Fig. 1f). The overall class distribution nevertheless differed between sample types (Fisher’s exact test, *p* = 5×10^-4^). Media DAPs were enriched for classically secreted proteins relative to lysate (14.0% versus 9.1%) and depleted of membrane proteins lacking a signal peptide (6.9% versus 11.6%), while shed and ectodomain proteins and GPI-anchored proportions were comparable (Fig. 1f).

Together, these results show that patient cerebral organoids carry a reproducible AD proteomic signature that is detectable across three independent platforms in both the conditioned media and the lysate. The media and lysate capture distinct fractions of this signature, the media enriched for secreted proteins and the lysate for intracellular and membrane proteins.

### Patient cerebral organoids detect nearly all literature-nominated AD biomarkers and drug targets

To benchmark the organoid model against the published AD proteomic literature, we systematically curated proteomic studies of patient plasma, CSF, and frontal cortex. Of 740 studies screened, 63 plasma, 42 CSF, and 16 frontal cortex studies met our inclusion criteria (Supplementary Fig. 3-c, Supplementary References). Together, these studies nominated 306 plasma biomarkers, 181 CSF biomarkers, and 123 cortex-derived drug targets (Supplementary Table 2, Fig. 2a). Seven proteins were shared across all three sample types (APOE, BDNF, GFAP, HGF, HP, SMOC1, and SPP1), with a further 38 shared between plasma and CSF (Supplementary Table 2, Fig. 2a). Notably, every included cortex study used MS alone, identifying a field-wide gap in the affinity-based proteomic interrogation of postmortem AD brain tissue.

**Figure 2.**
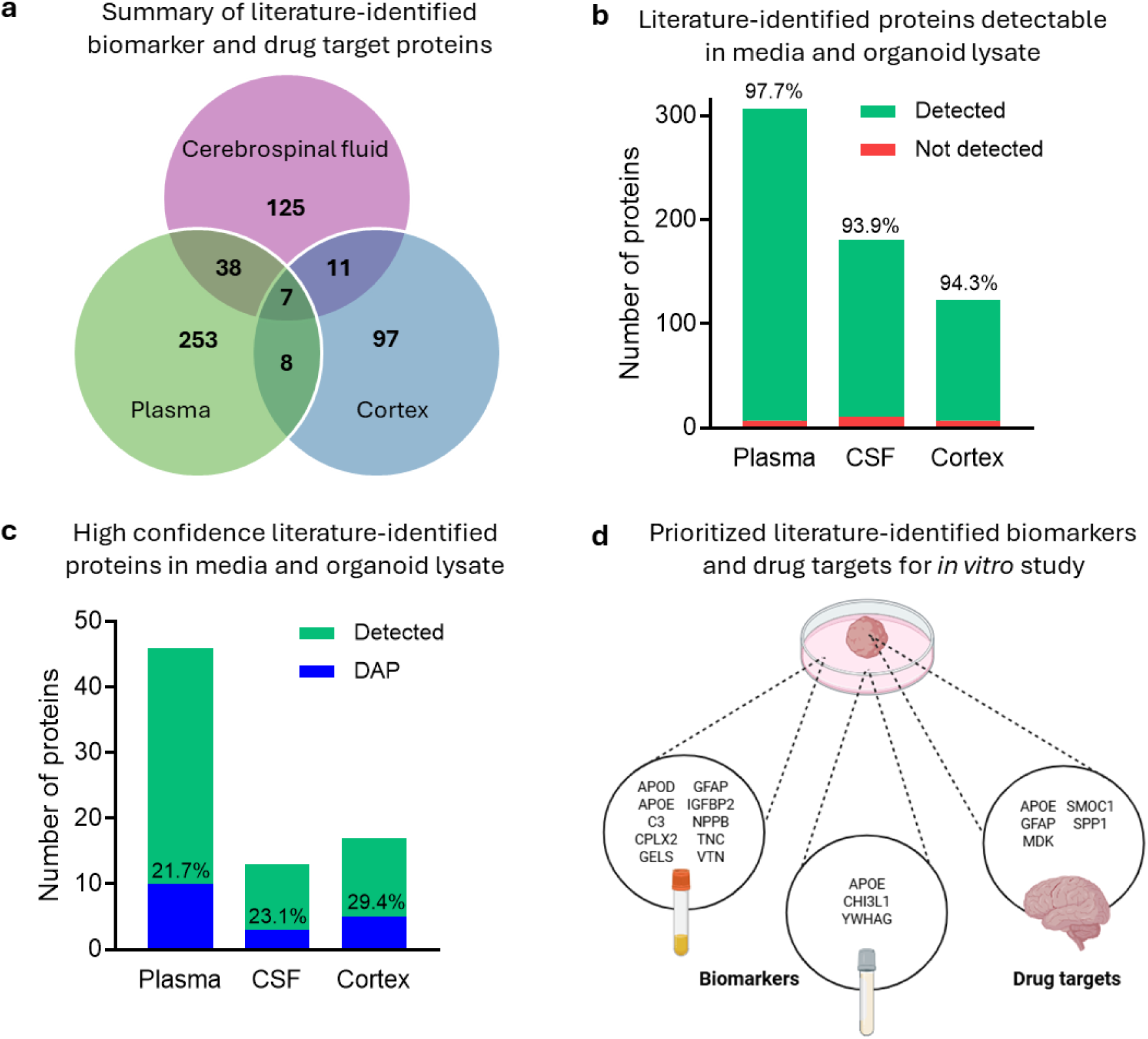
Patient cerebral organoids detect nearly all literature-nominated AD biomarkers and drug targets. (**a**) Venn diagram of AD biomarkers and drug targets nominated by the systematic review of patient plasma (306 proteins), CSF (181 proteins), and frontal cortex (123 proteins) studies. Seven proteins (APOE, BDNF, GFAP, HGF, HP, SMOC1, and SPP1) were nominated across all three sample types, with a further 38 shared between plasma and CSF. (**b**) Proportion of literature-nominated proteins detected in the matched organoid fraction. The conditioned media recovered 299 of 306 plasma biomarkers (97.7%) and 170 of 181 CSF biomarkers (93.9%), and the organoid lysate recovered 116 of 123 cortex-derived drug targets (94.3%). (**c**) Proportion of the most replicated literature candidates that were differentially abundant between AD and control organoids. DAPs accounted for 21.7% of plasma biomarkers (10/46), 23.1% of CSF biomarkers (3/13), and 29.4% of cortex drug targets (5/17). (**d**) Prioritized translational panel of convergent proteins that were both replicated in the literature and differentially abundant in the organoid model, comprising 12 candidate biomarkers in the conditioned media (APOD, APOE, C3, CPLX2, GELS, GFAP, IGFBP2, NPPB, TNC, VTN, CHI3L1, and YWHAG) and five candidate drug targets in the lysate (APOE, GFAP, MDK, SMOC1, and SPP1). Abbreviations: AD: Alzheimer’s disease; CSF: cerebrospinal fluid; DAP: differentially abundant protein.

The majority of these literature-nominated proteins were detectable in the organoid model. The conditioned media recovered 299 of 306 plasma biomarkers (97.7%) and 170 of 181 CSF biomarkers (93.9%), while the organoid lysate recovered 116 of 123 cortex-derived drug targets (94.3%) (Fig. 2b). The few undetected proteins were almost exclusively those with extremely low or absent brain expression in the Human Protein Atlas ^28^ and PaxDb ^29^, including lymphoid-restricted immunoglobulin chains and proteins absent from brain such as APOC4, ZNF587B, and ST8SIA5. This indicates that the detection gaps reflect tissue-of-origin biology rather than a limitation of the model.

We next restricted our analysis to the most heavily replicated candidates, namely plasma proteins reported in more than three studies (n = 46), CSF proteins reported in more than three studies (n = 13), and cortex proteins reported in more than two studies (n = 17). Every protein in each replicated set was measurable in the matched organoid fraction. A substantial subset was also differentially abundant between AD and control organoids, including 21.7% of plasma biomarkers (10/46), 23.1% of CSF biomarkers (3/13), and 29.4% of cortex drug targets (5/17) (Supplementary Table 2, Fig. 2c). These convergent proteins define a prioritized translational panel of 12 candidate biomarkers in the media (APOD, APOE, C3, CPLX2, GELS, GFAP, IGFBP2, NPPB, TNC, VTN, CHI3L1, and YWHAG) and five candidate drug targets in the lysate (APOE, GFAP, MDK, SMOC1, and SPP1) (Fig. 2d).

These data show that patient iPSC-derived cerebral organoids capture nearly all literature-nominated AD biomarkers and drug targets and reproduce the AD-associated change in approximately one in four of the most widely replicated candidates, including well-established AD proteins such as GFAP, SMOC1, and SPP1.

### Patient cerebral organoids reproduce the AD signatures of large clinical cohorts and nominate prioritized biomarkers and drug targets

To benchmark the organoid model against harmonized clinical cohort data, we leveraged AD multi-platform proteomic datasets from plasma (n = 13,623 individuals), CSF (n = 2,110 individuals), and dorsolateral prefrontal cortex (dlPFC, n = 1,275 donors) (Supplementary Table 3). These cohorts draw on genetically, racially, and clinically diverse populations sampled across more than twenty clinical sites, providing the independent, real-world comparison. At FDR < 0.05, the plasma cohorts yielded 139 DAPs by MS, 4,153 by SomaScan, and 342 by Olink (Supplementary Table 3; Supplementary Fig. 4a-c). The CSF cohorts yielded 490 by MS, 1,227 by SomaScan, and 394 by Olink (Supplementary Table 3; Supplementary Fig. 4d-f). The dlPFC yielded 4,871 by MS, with no postmortem brain datasets available from the affinity platforms (Supplementary Table 3). The organoid model again detected almost all these clinical DAPs, recovering 98.8% of plasma and 96.4% of CSF DAPs in the conditioned media and 96.1% of dlPFC DAPs in the lysate (Supplementary Table 3, Fig. 3a).

**Figure 3.**
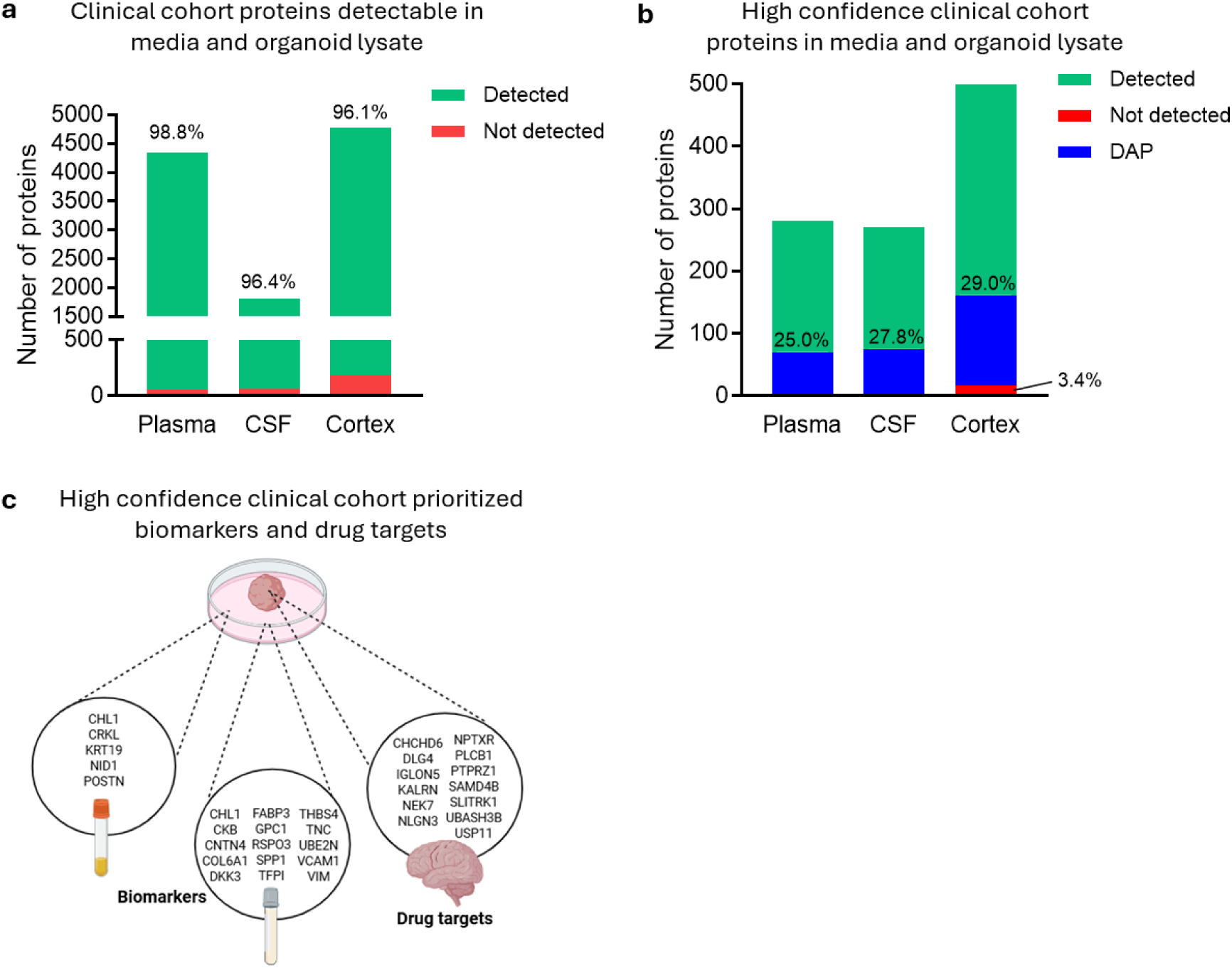
Patient cerebral organoids reproduce the AD signatures of large clinical cohorts and nominate prioritized biomarkers and drug targets. (**a**) Proportion of clinical cohort DAPs (FDR < 0.05) detected in the matched organoid fraction. The conditioned media recovered 98.8% of plasma and 96.4% of CSF DAPs, and the organoid lysate recovered 96.1% of dlPFC DAPs, drawn from cohorts of 13,623 plasma, 2,110 CSF, and 1,275 dlPFC samples. (**b**) Proportion of the highest confidence cohort DAPs that were also differentially abundant in the organoid. Of these, 70 of 280 plasma DAPs (25.0%), 75 of 270 CSF DAPs (27.8%), and 145 of 500 dlPFC DAPs (29.0%) reached DAP status in the organoid. (**c**) Prioritized proteins that were multi-platform DAPs in both the organoid and the human cohorts. Abbreviations: AD: Alzheimer’s disease; CSF: cerebrospinal fluid; DAP: differentially abundant protein; dlPFC: dorsolateral prefrontal cortex; MS: mass spectrometry.

We then focused on the highest confidence cohort DAPs, defined for plasma and CSF as those identified by two or more platforms, and for dlPFC as the top 500 MS DAPs (FDR < 1×10^-11^). Among the 280 plasma DAPs meeting these criteria, all were detectable in the conditioned media and 70 (25.0%) were themselves DAPs in the organoid. The 270 CSF DAPs identified by two or more platforms showed the same pattern, with all detectable in the media and 75 (27.8%) reaching DAP status, and of the top 500 dlPFC DAPs, 145 (29.0%) were also DAPs in the organoid lysate (Supplementary Table 3, Fig. 3b). The organoid therefore reproduced the AD-associated change in roughly one in four of the most reproducible clinical candidates, irrespective of sample type or platform.

To identify the proteins most amenable to translational study in patient organoids, we extracted those that were multi-platform DAPs in both the organoid and the human cohorts. Five proteins met this criterion across the plasma cohorts and conditioned media (CHL1, CRKL, KRT19, NID1, POSTN), and 15 across the CSF cohorts and media (CHL1, CKB, CNTN4, COL6A1, DKK3, FABP3, GPC1, RSPO3, SPP1, TFPI, THBS4, TNC, UBE2N, VCAM1, VIM) (Supplementary Table 3, Fig. 3c). At the tissue level, 194 proteins were multi-platform DAPs in the organoid lysate and DAPs in the dlPFC. We further prioritized these by requiring a concordant direction of change between the lysate and dlPFC, together with preferential brain expression, yielding 13 high-priority drug targets (CHCHD6, DLG4, IGLON5, KALRN, NEK7, NLGN3, NPTXR, PLCB1, PTPRZ1, SAMD4B, SLITRK1, UBASH3B, and USP11) (Supplementary Table 3, Fig. 3c, d). Of these, seven function in synaptic biology (DLG4, NLGN3, NPTXR, SLITRK1, IGLON5, KALRN, PLCB1), two in receptor tyrosine kinase signalling and protein turnover (UBASH3B, USP11), and the remaining four span mitochondrial integrity (CHCHD6), CNS developmental signalling (PTPRZ1), kinase activity (NEK7), and neuronal RNA regulation (SAMD4B) (Fig. 3d, Table 1).

**Table 1.**
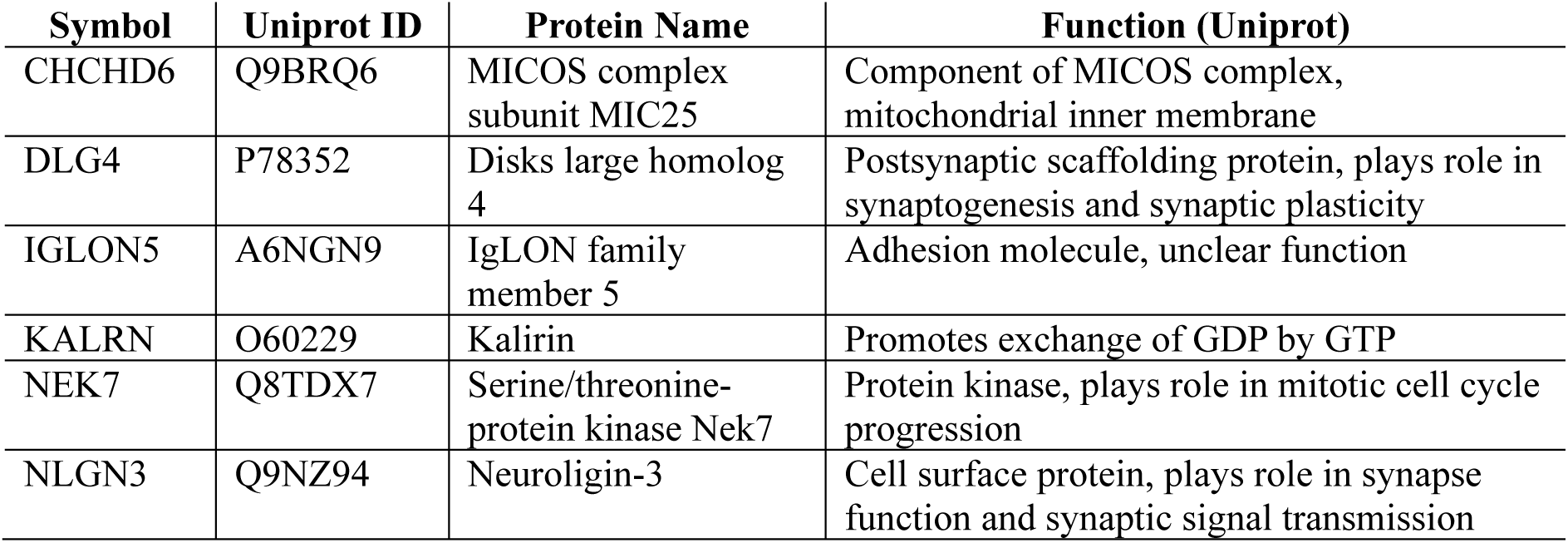
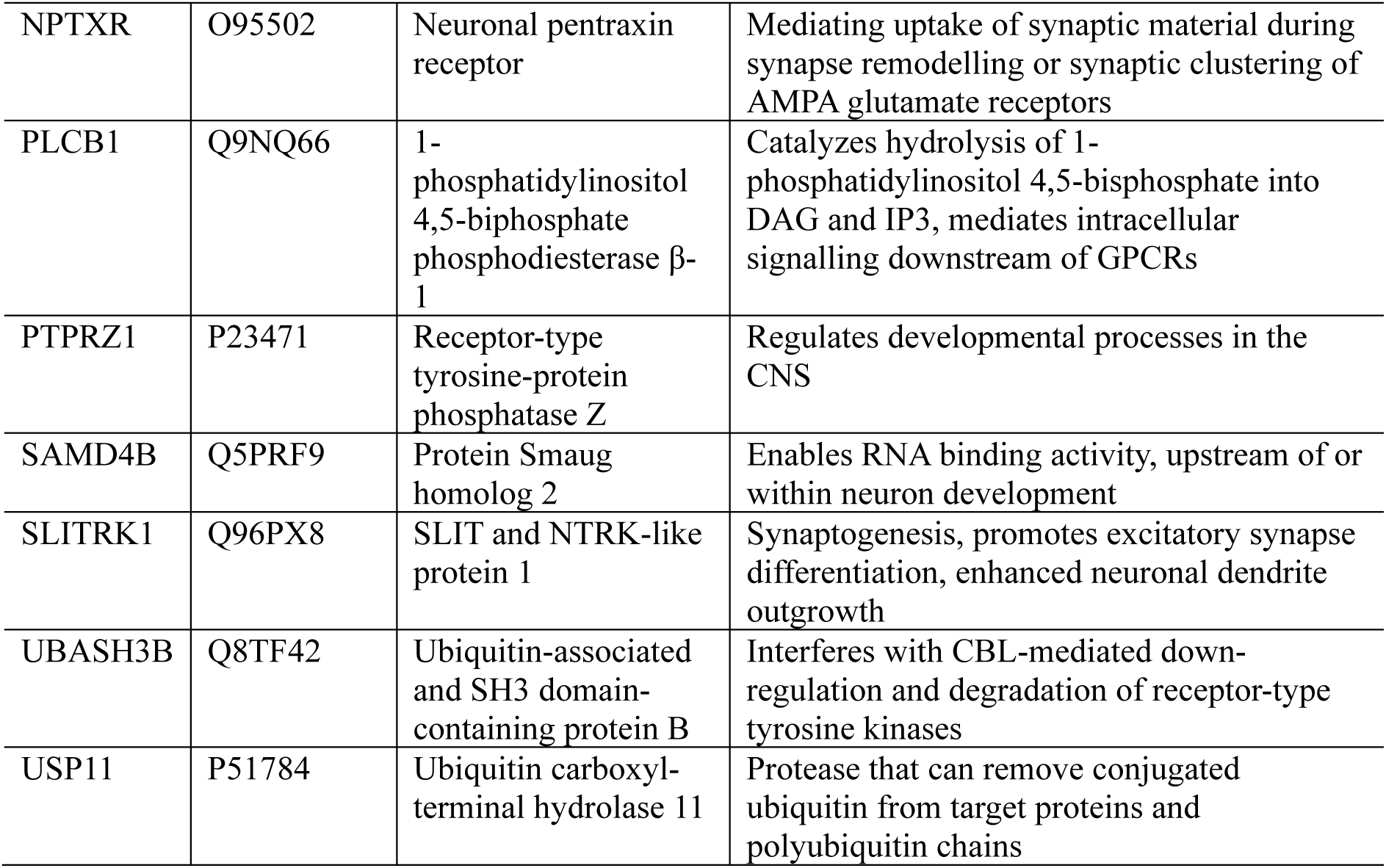
Biological function of the 13 prioritized AD drug targets.

Together, these findings show that patient cerebral organoids reproduce the disease-associated changes found across more than 17,000 clinical plasma, CSF, and cortex samples, recovering roughly one in four of the most reproducible cohort DAPs in each. This held across genetically and clinically diverse populations, indicating that the organoid model generalizes beyond the donors profiled here. From these convergent proteins, we nominate a prioritized panel of 5 plasma and 15 CSF biomarkers in the conditioned media and 13 drug targets in the lysate. These candidates are concentrated in the synaptic, mitochondrial, and proteostatic biology of AD and are immediately testable in the patient-derived model.

### Patient cerebral organoids reproduce the dysregulated biological processes of human AD

Recapitulation of individual proteins does not guarantee that an organoid reproduces the biological processes that define AD. We therefore asked whether the DAPs in patient organoids and in the human cohorts converge on the same dysregulated processes. For each platform, we compared the biological processes enriched among DAPs in the conditioned media against those in plasma and CSF, and in the organoid lysate against those in the dlPFC (Supplementary Table 4). Convergence between the conditioned media and biofluids tracked the depth of each platform. SomaScan recovered 116 enriched processes in the media, of which 36 were shared with CSF and 44 with plasma (Supplementary Table 4, Fig. 4a-c). The media-to-CSF overlap was again centered on neuronal and synaptic biology, with shared processes including axonogenesis, axon guidance, neuron differentiation, neuron projection development, and synaptic transmission. Olink recovered 43 enriched processes in the media and shared 10 with plasma, distributed across cell cycle regulation and checkpoints, the p53-mediated DNA damage response, and the cellular stress response. There were no enriched processes in the Olink CSF data, as more than 85% of the proteins on the panel were themselves DAPs. MS recovered no surviving enrichments in the media or plasma, consistent with the dynamic range limitations of MS in biofluids and low protein samples relative to the affinity platforms ^25,27^.

**Figure 4.**
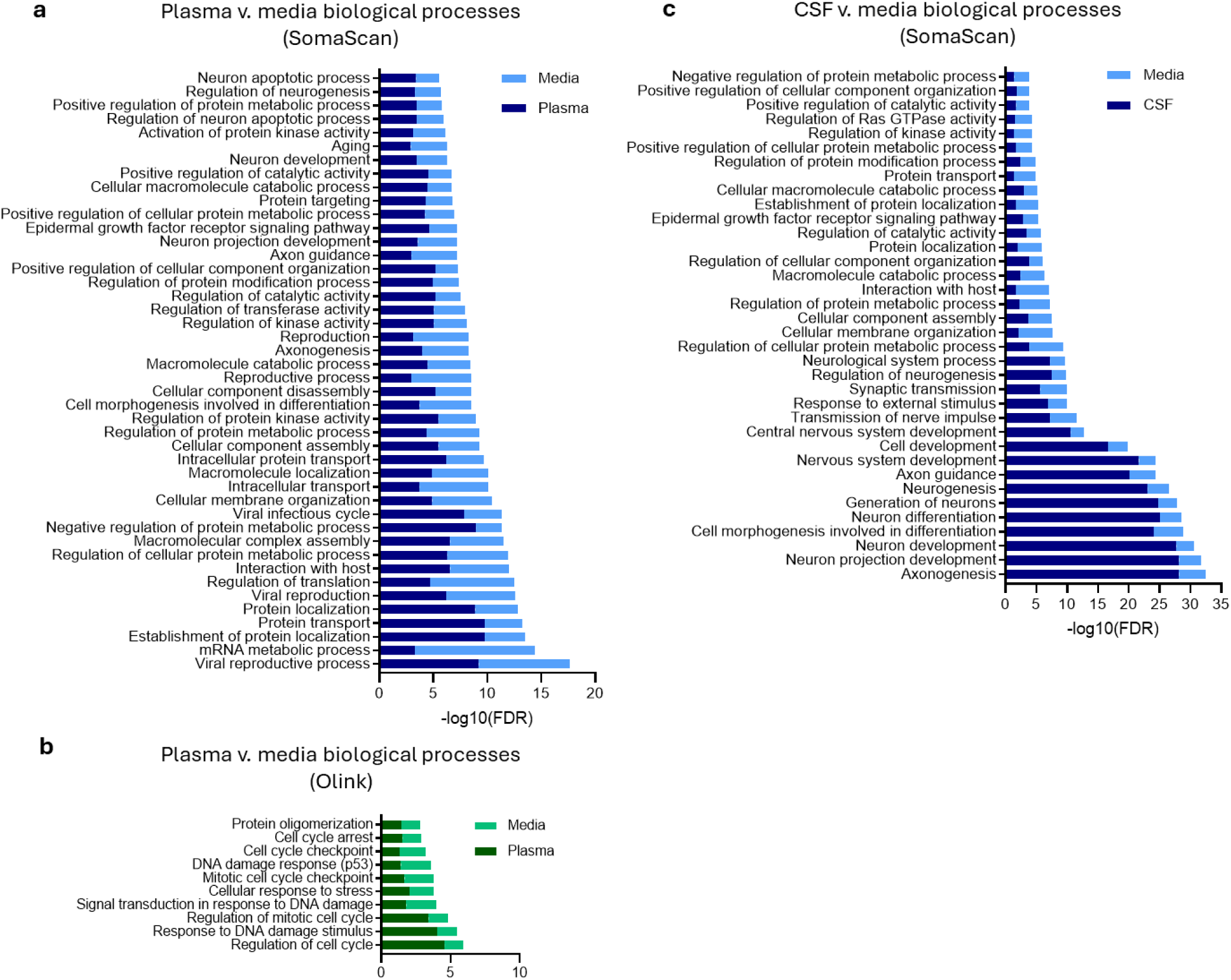
Patient cerebral organoid conditioned media reproduces the dysregulated biological processes of patient plasma and CSF. Comparison of biological processes (Gene Ontology; FDR < 0.05) enriched among biofluid and conditioned media DAPs. Biological processes enriched in plasma and conditioned media across (**a**) SomaScan and (**b**) Olink DAPs. (**c**) Biological processes enriched in CSF and conditioned media across SomaScan DAPs. Abbreviations: AD: Alzheimer’s disease; CSF: cerebrospinal fluid; DAP: differentially abundant protein.

The organoid lysate and human dlPFC shared 37 enriched processes by MS, 31 by SomaScan, and 34 by Olink (Supplementary Table 4, Fig. 5a-c). Protein localization, macromolecule localization, and cellular component assembly were shared across all three platforms, identifying these as the most robust biological signatures recovered by the model. Neuronal and synaptic processes, including axonogenesis, axon guidance, neuron projection development, and neuron differentiation, were shared between the lysate and dlPFC on LC/MS and SomaScan. Olink additionally recovered mitochondrial membrane organization, cytoskeleton organization, actin filament-based processes, and macromolecular complex assembly. The dlPFC was led by protein localization and transport, consistent with the proteostatic decline of the aged, end-stage AD brain ^13,30^. The organoid lysate was instead led by neurodevelopmental and morphogenic processes, consistent with the younger, fetal-like cellular state of cerebral organoids ^31–33^.

**Figure 4.**
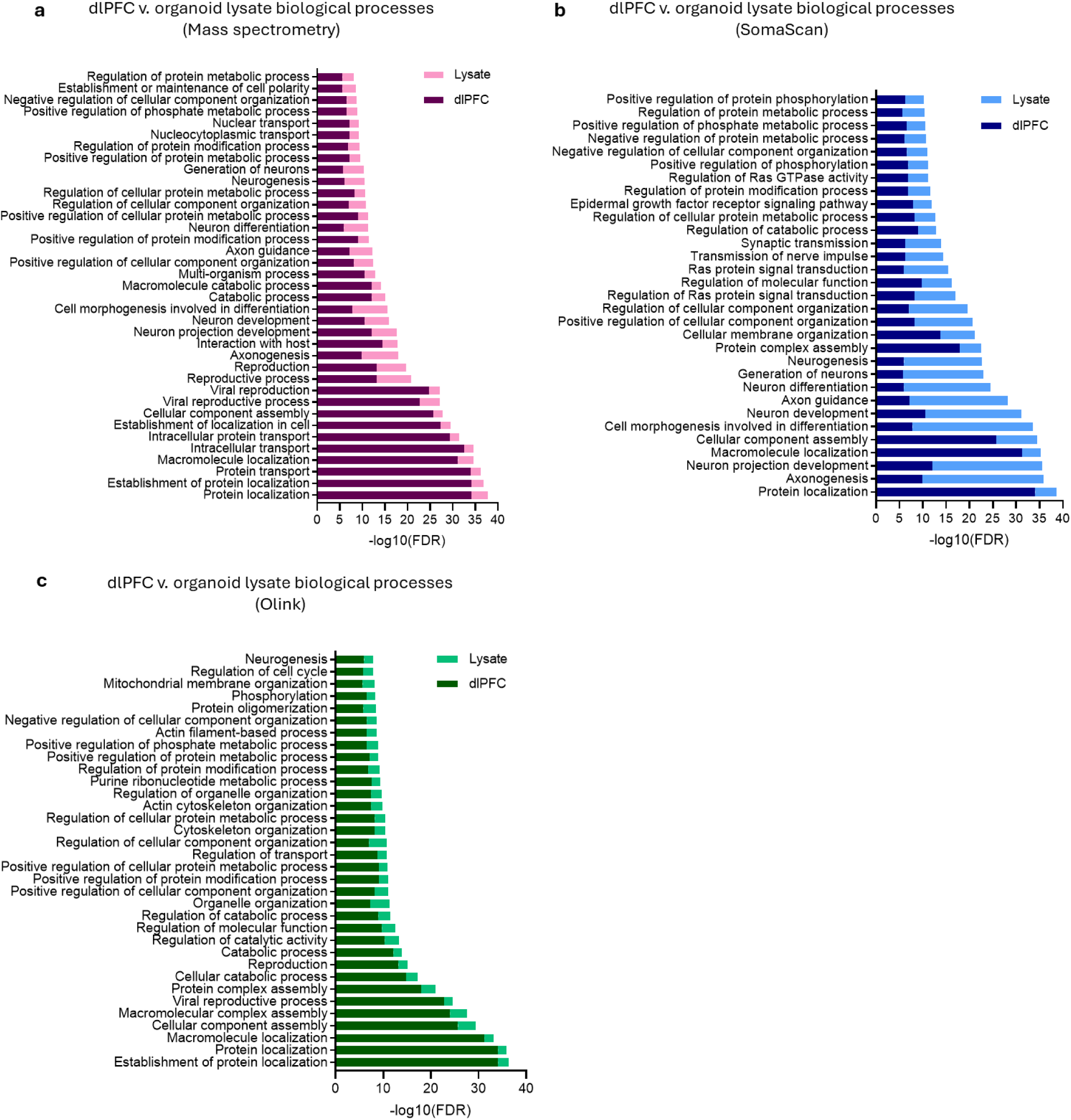
Patient cerebral organoid lysate reproduces the dysregulated biological processes of patient cortex. Comparison of biological processes (Gene Ontology; FDR < 0.05) enriched across the cortex and organoid lysate DAPs across (**a**) LC/MS, (**b**) SomaScan, and (**c**) Olink.

Taken together, these data show that patient cerebral organoids reproduce the dysregulated biology of human AD across cortex, CSF, and plasma, not only its individual proteins. The organoid lysate recovers the neuronal, cytoskeletal, and proteostatic processes of human cortex. The conditioned media recovers the neuronal and synaptic processes of patient CSF, along with the cell cycle and DNA damage response processes of patient plasma. The biological processes that define AD in the brain and biofluids can therefore be studied directly in a patient-derived model.

## DISCUSSION

Patient iPSC-derived cerebral organoids have become a leading human model for studying AD, yet their proteome has never been benchmarked against human disease. Here, we profiled the proteome of patient cerebral organoids on the three platforms that drive clinical AD discovery, LC/MS, SomaScan, and Olink, across both the organoid lysate and conditioned media. We compared this proteome against AD biomarkers and drug targets nominated by more than 120 published proteomic studies and against differentially abundant proteins from clinical cohorts spanning more than 17,000 plasma, CSF, and cortex samples. We found that patient cerebral organoids detect almost all clinically nominated candidates and reproduce the disease-associated change in roughly one in four of the most reproducible. This convergence held across plasma, CSF, and cortex, and extended to the synaptic, mitochondrial, and proteostatic biology of the disease. To our knowledge, this is the first multi-platform reference proteome of a patient-derived AD model, providing the field with a resource for advancing clinically nominated AD biomarkers and drug targets into pre-clinical research.

Against both the published literature and the clinical cohorts, patient cerebral organoids had near complete detectability of nominated AD proteins. From the literature, the conditioned media recovered 97.7% of nominated plasma biomarkers and 93.9% of CSF biomarkers, and the organoid lysate recovered 94.3% of cortex-derived drug targets. The same held against the clinical cohorts, where the media detected 98.8% of plasma and 96.4% of CSF DAPs and the lysate detected 96.1% of those from cortex. The few undetected proteins were largely lymphoid and peripheral, including immunoglobulin chains and proteins with little or no brain expression such as APOC4 and ST8SIA5 ^28,29^, reflecting the cellular makeup of the organoid rather than the depth of the platforms. Almost every AD biomarker and drug target nominated in these clinical cohorts is therefore expressed in a simplified patient organoid of neurons, astrocytes, and microglia, supporting its use as a translational model of AD.

Beyond detection, the organoid reproduced the disease-associated change in a substantial subset of these candidates. Roughly one in four of the most widely replicated literature candidates and the most reproducible cohort DAPs were themselves DAPs in the cerebral organoid, and this held across plasma, CSF, and cortex. These included several of the most reproducibly reported proteins in AD. Among the prioritized secretome biomarkers were the astrocyte-related proteins GFAP, which tracks astrocyte reactivity and rises in blood before cognitive decline ^34^ and CHI3L1, which is regulated by the astrocytes and modulates neuroinflammation ^35^. Among the prioritized lysate drug targets were SMOC1, which co-localizes with amyloid plaques ^36^ and rises in CSF nearly three decades before symptom onset in autosomal dominant AD ^37^, and SPP1, which mediates microglial engulfment of synapses ^38^. Each of these was differentially abundant between AD and control in the organoid, meaning the model captures not only the presence of these candidates but their dysregulation in disease.

We identified 13 prioritized drug targets in the organoid lysate, with roles in the synaptic, mitochondrial, and proteostatic processes affected in AD ^39–41^. Seven of these proteins play a role in synaptic structure and function. DLG4, which encodes PSD-95, is linked to synaptic change in AD ^42^ and protects synapses from amyloid-β ^43^. NPTXR, a presynaptic regulator of AMPA receptors, predicts clinical progression when measured in CSF ^44^. NLGN3 belongs to the neuroligin family of trans-synaptic adhesion molecules whose CSF concentrations are linked to neurodegeneration ^45^, and KALRN regulates dendritic spine integrity and has been implicated in synaptopathies ^46,47^. The remaining targets span other AD-relevant biology. CHCHD6 is a MICOS complex subunit whose loss drives amyloidogenic APP processing in human brain ^48^. NEK7 is a kinase required for NLRP3 inflammasome activation in AD neuroinflammation ^49,50^. USP11 is an X-linked deubiquitinase and has been linked to tauopathy and AD in women ^51,52^. Each of these 13 targets was differentially abundant in both human AD cortex and the patient organoid, identifying a set of drug targets that are altered in the human disease and immediately directly testable in a patient-derived model.

The organoid and human cohorts also converged on biological processes. The conditioned media tracked patient CSF on synaptic and axonal biology, both of which are early and central features of AD ^53,54^. It also aligned with patient plasma on cell cycle regulation and the p53-mediated DNA damage response, processes that are aberrantly activated in AD neurons ^55–58^. Organoid lysate and human cortex shared dozens of enriched processes across platforms, all of which have been supported by previous AD findings, including protein and macromolecule localization ^13^, neuronal and axonal development ^53,54^, and cytoskeletal ^59^ and mitochondrial ^60^ organization. They also differed, however, in their leading processes. The cortex was led by protein transport and proteostatic processes, consistent with the proteostatic decline of the aged, end-stage postmortem brain ^13,30^, whereas the organoid was led by neurodevelopmental and morphogenic processes. Cerebral organoids resemble fetal and early postnatal brain ^31–33^, and the cells profiled here likely sit at an earlier point on the disease trajectory than postmortem cortex. The organoid therefore reproduces the process-level biology of human AD, with the difference in leading processes reflecting its younger developmental stage rather than a divergence from the disease.

Our study has several limitations. First, we profiled organoids from one AD and one control donor. It is often held that three to five iPSC lines are needed to account for donor genetic and phenotypic heterogeneity ^61,62^, however we note that this arbitrary threshold has no empirical basis. Reliably accounting for heterogeneity can require upward of 50 donors, which no iPSC disease modelling study can reach ^63^. Generalizability here comes not from donor number but from convergence with more than 17,000 genetically and clinically diverse individuals across the clinical cohorts. Further, our approach is in line with other recent work showing that iPSC models derived from a single donor are informative when used to confirm proteomic findings from large-scale AD clinical cohorts ^64^. Second, cerebral organoids have limited cellular diversity and structural complexity. Although we integrate in iPSC-derived microglia, moving beyond the traditional neuron-astrocyte composition ^65^, they lack vasculature, a blood-brain barrier, oligodendrocytes, and peripheral immune cells that contribute to pathophysiology *in vivo* ^66^. Protocols do exist to engineer vasculature and a blood-brain barrier ^67,68^, oligodendrocytes ^69^, and peripheral immune cells ^70,71^ are available or emerging, but are less established and add considerable technical complexity. We therefore benchmarked a widely used cerebral organoid model, and extending this benchmark to more complex models is a clear next step. Third, the depth of the conditioned media proteome by MS is limited and could have been increased by depleting high-abundance proteins or by culturing in low-protein media. However, depletion can remove genuine protein biomarkers ^72,73^ and low-protein media compromises organoid health and would likely increase stress-related proteins. The media proteome therefore reflects organoid biology rather than sample handling, and the two affinity platforms recovered secreted proteins beyond the reach of MS.

Together, our findings establish the patient cerebral organoid as a translationally relevant model of AD and provide a reference proteome that spans the three proteomic platforms driving clinical discovery. Clinical cohorts now nominate biomarkers and drug targets faster than they can be tested in models of the human disease. By defining which candidates are reproduced in a patient organoid, this resource connects cohort-scale discovery to pre-clinical investigation and provides a foundation for the mechanistic and therapeutic study of AD in a patient-derived system.

## Supporting information

Supplementary References

## RESOURCE AVAILABILITY

### Lead Contact

Caitlin A. Finney; 176 Hawkesbury Road, Westmead, New South Wales, Australia 2145

### Materials availability Data and code availability

All the proteomic data generated in this study are available in Supplementary Tables 1 and 3.

The GNPC data used in the current study is available by request via the AD Discovery Portal (https://discover.alzheimersdata.org/). Olink PEA plasma data from the Emory Goizueta ADRC Multi-Platform Study is available through the Synapse database (https://www.synapse.org/Synapse:syn30549757/). CSF mass spectrometry data from the Emory 300 study is available through the Synapse database (https://www.synapse.org/Synapse:syn68293899). CSF Olink PEA data from the PRIDE DLB study is available through the Synapse database (https://www.synapse.org/Synapse:syn52282088/). Researchers who wish to access these controlled datasets are required to submit a Data Use Agreement. More information can be found at https://discover.alzheimersdata.org/ and https://adknowledgeportal.synapse.org/Data%20Access.

## ACKNOWLEDGEMENTS

This work was supported by the Australian Government’s Medical Research Future Fund MRF2040081 (C.A.F. & A.S.), MRF2052401 (A.S. & C.A.F.); James N. Kirby Foundation (C.A.F.); Dementia Australia Research Foundation Bondi2BlueMtns Project Grant (C.A.F, H.M.W. & R.H.S.); the NIH P30AG072973 (R.H.S., J.M.B., H.M.W., C.S.); the Victorian State Government Operational Infrastructure Support Program (S.W.Z.O.; R.B.); Australian Government NHMRC IRIIS to the Walter and Eliza Hall Institute of Medical Research (S.W.Z.O.; R.B.); Australian Government Research Training Program (RTP) scholarship (T.C.Y.T.); Australian Academy of Technological Sciences & Engineering Elevate Postgraduate Research Scholarship (T.C.Y.T.); Children’s Miracle Network (K.W.); and philanthropic funding from the John & Anne Leece Family (C.A.F. & A.S.), Paul & Valeria Ainsworth Family (C.A.F.), and Neil and Norma Hill Foundation (C.A.F).

The authors wish to thank the cohort contributors, participants, donors, and their families who helped make this research possible. The results published here are in whole or in part based on data obtained from the AD Knowledge Portal (https://adknowledgeportal.org/). Data generation was supported by the following NIH grants: U01AG046139, U01AG046170, U01AG061357, U01AG061356, U01AG061359, and R01AG067025. We thank the participants of participants of the Religious Order Study, Memory and Aging Project, the Minority Aging Research Study, Rush Alzheimer’s Disease Research Center, Mount Sinai/JJ Peters VA Medical Center NIH Brain and Tissue Repository, National Institute of Mental Health Human Brain Collection Core (NIMH HBCC), Mayo Clinic Brain Bank, Sun Health Research Institute Brain and Body Donation Program, Goizueta Alzheimer’s Disease Research Center, New York Brain Bank at Columbia University, New York Genome Center and the Biggs Institute Brain Bank for their generous donations. Data and analysis contributing investigators include Nilüfer Ertekin-Taner, Minerva Carrasquillo, Mariet Allen (Mayo Clinic, Jacksonville, FL), David Bennett, Lisa Barnes (Rush University), Philip De Jager, Vilas Menon (Columbia University), Bin Zhang, Vahram Haroutanian (Icahn School of Medicine at Mount Sinai), Allan Levey, Nick Seyfried (Emory University), Rima Kaddurah-Daouk (Duke University), Steve Finkbeiner (University of California-San Francisco/Gladstone Institutes), Daifeng Wang (University of Wisconsin-Madison), Stefano Marenco (NIMH HBCC), Anna Greenwood, Abby Vander Linden, Laura Heath, William Poehlman (Sage Bionetworks).

We acknowledge the support of the Children’s Medical Research Institute Biomedical Proteomics Facility and use of equipment funded by the Sargents Charitable Foundation.

## AUTHOR CONTRIBUTIONS

Conceptualization, Data curation, Formal analysis, Funding acquisition: A.S. & C.A.F.; Investigation, Writing – review & editing: S.T., C.L., S.W., T.C.Y.T., M.E.G., S.W.Z.O., R.B., C.S., J.M.B., K.A.W., R.H.S., H.M.W., A.S., C.A.F.; Methodology: S.T., M.E.G., S.W.Z.O., R.B., A.S., C.A.F; Writing – original draft: S.T., C.A.F.; Software: A.S.; Project administration, Resources, Supervision: C.A.F.

## METHODS

### Patient iPSC-derived cerebral organoids

#### Induced pluripotent cell derivation

Post-mortem dermal fibroblast samples were collected from two female donors, one healthy control aged 72 and AD aged 84, from the University of Kansas Alzheimer’s Disease Research Center cohort. Both donors provided written consent to donation upon death under approval by the University of Kansas Medical Center’s Institutional Review Board (FWA00003411). iPSCs were generated from dermal fibroblasts using a CytoTune-iPSC 2.0 Sendai Reprogramming kit (ThermoFisher Scientific, A16517). Six-well plate were coated with Matrigel (0.08mg/ml; Corning, 356234) diluted in DPBS prior to cell seeding. iPSCs were maintained in StemFlex medium (Gibco, A3349401) containing 100U/ml penicillin-streptomycin (Gibco, 15140122), with media refreshed every 48 hours. Weekly passaging was performed using ReLeSR (StemCell Technologies, 15140122) according to the manufacturer’s protocol, and cultures were supplemented with 10μM Y-27632 ROCK inhibitor (StemCell Technologies, 72302) for the first 24 hours post-passage to promote cell survival.

#### iPSC-derived microglia differentiation

Microglia were generated from iPSCs through a stepwise differentiation protocol. iPSCs were first directed toward a hematopoietic fate using a STEMdiff Hematopoietic kit (StemCell Technologies, 05310), with cells seeded into Matrigel-coated 24-well plates (0.04mg/ml) in HPC Medium A. After 72 hours, medium was exchanged for HPC Medium B for a further nine days. Resulting HPCs were harvested and replated in STEMdiff Microglia Differentiation medium (StemCell Technologies, 100-0019), with half-medium changes performed every 48 hours over a 24-day period and a single passage carried out at day 12. Differentiated microglia were then passaged into STEMDiff Microglia Maturation Medium (StemCell Technologies, 100-0020) as per the manufacturer’s instructions. Fresh media was added every other day for a total of 10 days of maturation. Cells were then harvested using Accutase (ThermoFisher Scientific, 00-4555-56) prior to integration into embryoid bodies.

#### Embryoid bodies

iPSCs were passaged three times to stabilize the line. After wells reached 90 to 100% confluence, iPSCs were removed from the well using ReLeSR (StemCell Technologies, 15140122) and 1.5 x 10^5^ cells were added into each well of a 24-well plate. Plates were pre-treated with Anti-Adherence Rinsing Solution (StemCell Technologies, 07010), rinsed with PBS and prepared with 0.5ml/well of StemFlex medium (Gibco, A3349401) supplemented with 100U/ml penicillin-streptomycin and 10μM Y-27632 ROCK inhibitor (StemCell Technologies, 72302). Medium was changed every two days until iPSCs had formed large embryoid bodies (EBs) approximately 250μm in diameter.

#### Cerebral organoids

To create organoids, EBs were passaged into 24-well plates pre-rinsed with Anti-Adherence Rinsing Solution (StemCell Technologies, 07010) and PBS at approximately 1-2 EBs per well. Plates were prepared with 0.5ml/well N2/B27 week 1 medium (Table 2) supplemented with 10μM Y-27632 ROCK inhibitor (StemCell Technologies, 72302). Mature microglia were harvested and added to each well of EBs at a concentration of 0.8 x 10^5^ cells/well. Fresh N2/B27 medium was added to each well every 4 hours and organoids were passaged at the end of every stage in a four-stage protocol adapted from the Lancaster protocol ^65^ (Table 2). Briefly, organoids underwent initial differentiation to a neural progenitor cell (NPC) stage in base N2/B27 medium supplemented with dorsomorphin (StemCell Technologies, 72102), SB431542 (StemCell Technologies, 72234) and CHIR99021 (StemCell Technologies, 72054), and thiazovivin (StemCell Technologies, 72254) for one week (Methods Table 1). Media was then changed to N2/B27 medium supplemented with epidermal growth factor (EGF; StemCell Technologies, 78006) and basic fibroblast growth factor (bFGF; StemCell Technologies, 78003) for two weeks to promote neuronal and astrocytic differentiation (Table 2). Cerebral organoids were then passaged into N2/B27 medium supplemented with dibutyryl cyclic-AMP (dcAMP; StemCell Technologies, 73882), brain derived neurotrophic factor (BDNF; StemCell Technologies, 78005), glial cell line-derived neurotrophic factor (GDNF; StemCell Technologies, 78058), and sodium pyruvate (ThermoFisher Scientific, 11360070) for three weeks (Table 2). This was then replaced with growth factor-free N2/B27 medium for an additional two weeks. Cerebral organoids were the removed from the media, washed in DPBS three times for 10 minutes each, and flash frozen in liquid nitrogen prior to being stored at -80°C. A total of 1ml of conditioned media from each organoid well was also flash frozen in liquid nitrogen and stored at -80°C.

**Table 2.**
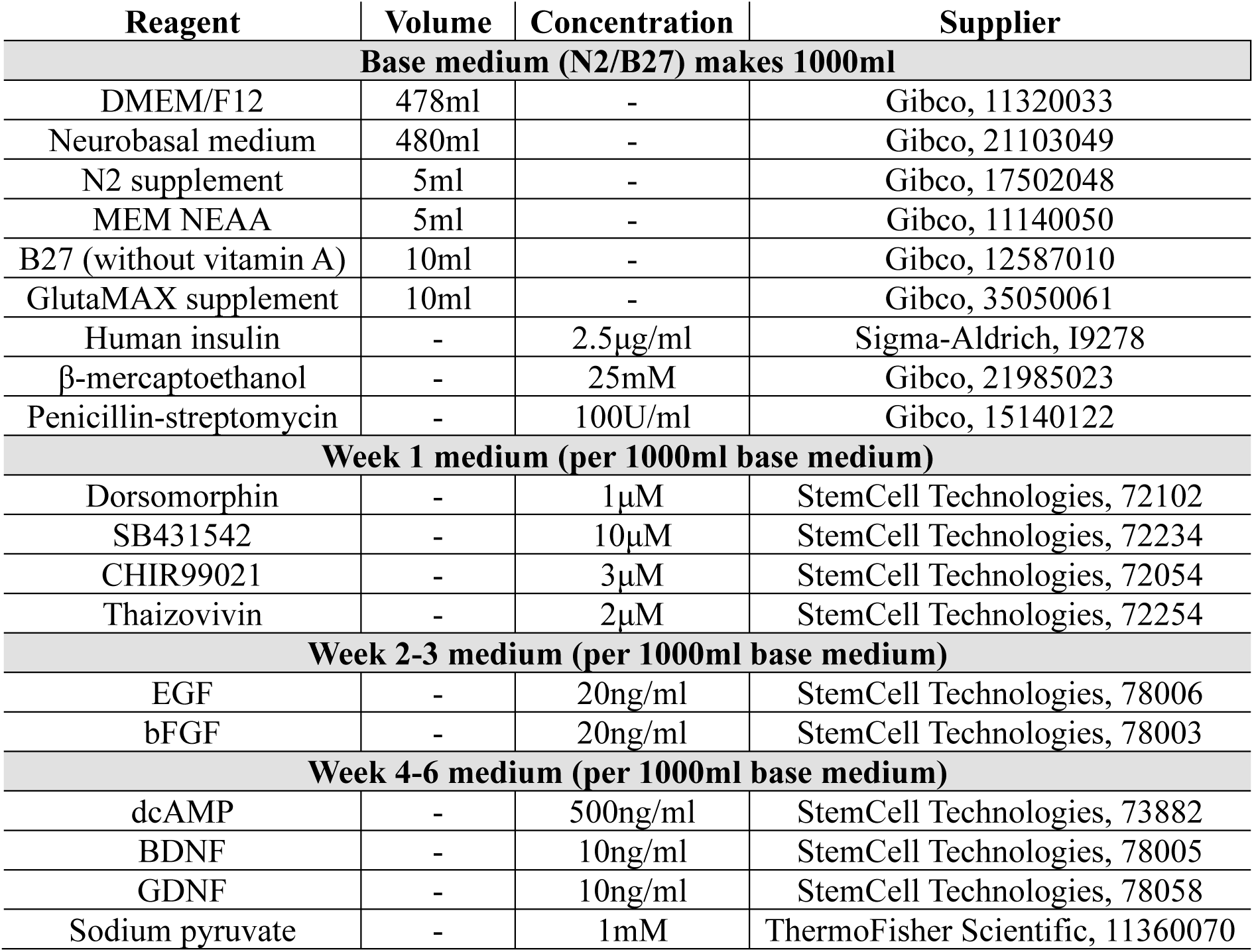
List of media reagents for cerebral organoid differentiation and maturation.

### Cerebral organoid proteomics

#### Label free mass spectrometry

Cerebral organoids and conditioned media were collected at 8 weeks, immediately snap frozen in liquid nitrogen, and stored at -20°C until further processing. For protein extraction, samples were resuspended in a lysis buffer consisting of 0.2% n-dodecyl-β-D-maltoside (DDM) in 50mM triethylammonium bicarbonate (TEAB) supplemented with cOmplete EDTA-free Protease Inhibitor Cocktail (added as per manufacturer’s instructions). Organoids were mechanically disrupted using a drill-mounted pestle for 10 seconds, then heated to 85°C for 10 minutes. Samples were cooled and incubated with 12.5U of benzonase for 30 min at 37°C. The samples were incubated with 10mM tris(2-carboxyethyl)phosphine hydrochloride (TCEP) for five min at 85°C, then cooled and incubated with 20mM iodoacetamide at 23°C for 30 min. Enzymatic digestion was initiated by the addition of 2µg trypsin (Sigma, Trypzean) at 42°C for two hours, followed by a supplementary 2µg trypsin at 33°C for a further six hours. Digestion was quenched by acidification with 0.5µl formic acid, after which samples were diluted in 30µl of 0.1% trifluoroacetic acid and desalted using the STAGE tip method ^74^.

Peptide samples were separated and analyzed by LC-MS/MS using a Vanquish Neo UHPLC system coupled to an Astral Orbitrap mass spectrometer (ThermoFisher Scientific). Samples were first trapped on a Pepmap Neo C18 trap column (5 µm particles, 300 µm i.d. x 5 mm; ThermoFisher Scientific) at a flow rate of up to 10µl/min and a maximum pressure of 800 bar, before online elution through a 15cm x 150µm i.d. Pepmap C18 EASY-Spray analytical column (2µm, 100 Å particles; ThermoFisher Scientific) heated to 40°C. Electrospray ionisation was performed at 1.9 kV, with an S-lens RF level of 40 and a capillary temperature of 280°C. Chromatographic separation employed a binary solvent system comprising buffer A (0.1% formic acid in water) and buffer B (0.1% formic acid in acetonitrile). Following sample loading at 3.6% buffer B, peptides were eluted at 1µl/min through a stepped gradient: 7.2-25.2% buffer B over 19.7 minutes, 25.2-31.5% buffer B over 3.7 minutes, 31.5-49.5% buffer B over 0.4 minutes, and finally ramped to 90% buffer B at 3µl/min over 0.5 minutes and held for 0.7 minutes, with a total MS acquisition window of 26 minutes. Full MS scans were acquired in the Orbitrap at a resolution of 240,000, with an AGC target of 5,000,000 and a maximum ion injection time of 3ms over a scan range of m/z 380-980. Data-independent acquisition (DIA) MS/MS scans were performed in the Astral analyser using 2m/z isolation windows, an automatic gain control (AGC) target of 50,000, a maximum ion injection time of 3ms, a scan range of m/z 150-2,000, and a normalised collision energy of 25. The cycle time was set to 0.6 seconds.

Raw data were processed using DIA-NN v2.2 ^75^. Peptide-spectrum matching was performed against an *in silico* spectral library generated from the canonical Human UniProt reference proteome (downloaded September 30 2025; 20,663 entries). Database search parameters included trypsin/P digestion with up to one missed cleavage, a peptide length range of 7-30 residues, and carbamidomethylation of cysteine as a fixed modification, with N-terminal methionine excision permitted. MS2 and MS1 mass tolerances were set to 10ppm and 4ppm respectively. Identifications were filtered to a precursor-level false discovery rate of 1%. Match-between-runs was enabled, heuristic protein inference was enabled, and all remaining parameters were left at DIA-NN v2.2 defaults.

#### Protein extraction

Organoids were weighed and an appropriate amount of Tissue Protein Extraction Reagent (T-PER) (ThermoFisher Scientific, 78510) was calculated at a 1:10 (w/v) ratio of tissue (mg) to reagent (μl) for SomaScan and 1:5 for Olink cerebral organoid samples. 100x HALT protease inhibitor cocktail (ThermoFisher Scientific, 88786) was added to T-PER at a concentration of 1:100μl for SomaScan organoid samples. Cerebral organoids were then homogenized using indirect sonication and centrifuged at 10,000 x g for five minutes at 4°C. Supernatant was then collected on ice and samples diluted to a concentration of 200μg/ml using PBS for SomaScan or 500μg/ml using T-PER for Olink. Conditioned media aliquots of 1ml as well as unconditioned (blank) base media were thawed and centrifuged at 10,000 x g for five minutes at 4°C. Supernatant was collected on ice.

#### SomaScan 11k assay

The SomaScan v5.0 assay (SomaLogic) was performed on conditioned media and cerebral organoid lysate samples. The assay detects approximately 11,000 proteins using slow off-rate modified aptamers (SOMAmers) that contain chemically modified nucleotides that bind with high specificity and affinity to target proteins ^76^. Protein measurements are provided in relative fluorescent units (RFU). Raw data was provided by SomaLogic following standardization and normalization.

#### Olink Explore HT assay

Proximity Extension Assay (PEA) proteomics was performed using Olink Explore-HT panel reagents (Olink Proteomics, P/N 98100). 300nL of sample was added to 600nL of multiplexed dual oligo-labelled detection antibodies in incubation solution, with all samples replicated across eight blocks of multiplexed assays comprising the complete Explore-HT panel. For all lysate and media samples, undiluted sample was transferred for each assay block, in place of the serial dilutions generally used for human plasma samples in blocks 5:8. After overnight incubation at 4°C, the PEA barcodes were extended, amplified and indexed with the following thermocycling parameters: 50°C for 20min; 95°C for 5min; 25x (95°C for 30sec; 54°C for 1min; 60°C for 1min). The products were then pooled and analyzed per block for expected size profile using Tapestation D1000 screentape (Agilent). DNA sequencing was then performed on Illumina NovaSeq X Plus with 10B flow cell. Reads were demultiplexed using ngs2counts (v6.2.0) and processed to NPX values by NPX Map software (v2.0.0, Olink Proteomics), by normalizing to the internal Extension Control and the external Plate Control ^77^.

### Clinical cohorts

#### Global Neurodegeneration Proteomics Consortium (GNPC) cohort

The GNPC cohort represents the largest aggregated proteomic resource for neurodegenerative diseases to date, drawing on samples from more than 20 clinical sites across the USA, UK, and Europe ^21^. The current study utilized plasmas and CSF samples from cognitively unimpaired controls (n = 9,059 plasma; n = 931 CSF) and individuals with clinically diagnosed AD (n = 4,476 plasma; n = 659 CSF; Supplementary Table 3). AD diagnosis was established at each contributed site according to previously described criteria ^21^, incorporating Clinical Dementia Rating (CDR) scores ≥ 1, Mini-Mental State Examination (MMSE) scores ≤ 24, and/or Montreal Cognitive Assessment (MoCA) scores ≤ 23. Demographic and clinical data, alongside biofluid samples, were obtained from a single study visit per participant (Supplementary Table 3). All participants provided written informed consent prior to enrolment, and ethical approval was granted by the relevant institutional review board at each site ^21^. Plasma proteomic data was generated using TMT mass spectrometry. Plasma and CSF proteomic data were generated using the SomaScan v4.1 assay (SomaLogic) ^21^. This aptamer-based assay enables the simultaneous measurement of approximately 7,000 proteins. Raw protein abundance, quantified using relative fluorescence units (RFU), were provided by SomaLogic following internal standardization, normalization, and calibration procedures, including adaptive normalization by maximum likelihood (ANML). All aptamers were mapped to UniProt identifiers. Full details of the dataset construction and harmonization across sites have been described previously ^21^.

#### Emory Goizueta Alzheimer’s Disease Research Center (ADRC) Multi-Platform study

The Emory Goizueta ADRC Multi-Platform study generated multi-platform proteomic data for plasma from n = 18 AD patients and n = 18 unimpaired controls (Supplementary Table 3). All participants provided written informed consent under ethical approval from the Emory University institutional review board ^22^. The plasma proteome was profiled using 13 human Olink Target 96 panels (cardiometabolic, cardiovascular II, cardiovascular III, cell regulation, development, immune response, inflammation, metabolism, neuro-exploratory, neurology, oncology II, oncology III, and organ damage) as described elsewhere ^22^.

#### Emory Goizueta ADRC 300 study

The Emory Goizueta ADRC 300 study generated TMT-MS proteomic data for CSF samples from n = 140 unimpaired controls and n = 160 AD patients (Supplementary Table 3) as described elsewhere ^78^. AD was diagnosed based on the NIA research framework ^79^. All participants provided written informed consent under ethical approval from the Emory University institutional review board ^78^.

#### PRIDE DLB study

The PRIDE DLB study draws on samples generated from the Amsterdam Dementia Cohort and DEvELOP ^80^ and included n = 110 unimpaired controls and n = 110 AD patients (Supplementary Table 3). AD patients were identified based on NINCDS-ADRDA criteria ^81^. A total of 979 proteins in the CSF were measured using 11 Olink Target Panels (cardiometabolic, cardiovascular II and III, cell regulation, development, immune response, inflammation, metabolism, neurology, oncology II, and organ damage), as described elsewhere ^80^. All participants provided written informed consent and the study was approved by the relevant institutional ethics review board at each clinical collection site ^80^.

#### Accelerating Medicines Partnership-AD (AMP-AD) Diverse Cohorts study

The AMP-AD Diverse Cohorts study is a cross-consortium initiative that generated harmonized, high-throughput multi-omic data from post-mortem brain tissue. AD cases were pathologically confirmed as described previously ^82^. Tissue samples from the dorsolateral prefrontal cortex (dlPFC) were sourced from four contributing institutions: Mayo Clinic, Rush University, Mount Sinai University Hospital, and Emory University. The present study incorporated dlPFC data from cognitively unimpaired controls (n = 399) and AD cases (n = 876). Full demographic information for the included donors is provided in Supplementary Table 3.

Written informed consent was obtained from all participants prior to donation, and all study protocols received ethical approval from the relevant institutional review boards ^82^. Proteomic profiling of the dlPFC homogenates was carried out by TMT mass spectrometry as previously described ^82^. To ensure data quality, only proteins with ≤ 30% missing values across the sample set were retained, with remaining missing values imputed using per-protein median values. Residual batch effects were corrected by fitting a linear regression model, consistent with the approach used in the original study ^82^.

### Statistical analyses

For LC/MS, DIA-NN protein-group intensities were log_2_-transformed and filtered within each matrix type to protein groups detected in at least 70% of samples. Protein groups were then mapped to their UniProt identifiers. Missing values (threshold <30%) were imputed using a MinProb strategy under a missing-not-at-random assumption, drawing replacement values from a normal distribution centred at the first percentile of observed values per protein with standard deviation equal to the observed standard deviation (tune factor of 1). Two sensitivity analyses were run in parallel, one using median imputation and one using complete case proteins only. Imputation sensitivity was assessed by Spearman correlation of fold changes between the MinProb and median approaches. SomaScan RFUs were log_2_-transformed and filtered to aptamers detected, defined as non-missing and non-zero, in at least 80% of samples. Near-zero-variance aptamers were removed. UniProt identifiers and gene symbols were taken from the analyte annotation. Olink measurements were filtered to protein assays from experimental samples that passed quality control, yielding 118,800 observations. Where multiple OlinkIDs mapped to the same UniProt accession within a sample, NPX values were averaged. Data were reshaped to a sample-by-protein matrix of 5,400 proteins, all detected in every sample. Differential abundance was assessed using limma with empirical Bayes moderation. *p* values were adjusted for multiple testing with the Benjamini-Hochberg procedure and proteins with an adjusted *p* value < 0.05 were considered differentially abundant (DAP).

Platform overlap was visualised using UpSet plots for both organoid and media matrices. Only proteins passing platform-specific quality filters were included. All analyses were performed in R (version 4.5.1). SomaScan files were parsed with ‘SomaDataIO’, differential expression was performed with ‘limma’, and data manipulation and visualisation used ‘matrixStats’, ‘dplyr’, ‘tidyr’, ‘ggplot2’, ‘ggrepel’, ‘pheatmap’, and ‘UpSetR’.

Protein-protein enrichment analyses were used to assess the biological function enriched by DAPs using NetworkAnalyst (v3.2) ^83–85^. Generic protein-protein interactions were identified using a zero-order network in the STRING database ^86^. Interactions were limited to those with a high confidence (>700 score) and with experimental evidence. The Gene Ontology database was used to identify biological processes and significance determined by FDR < 0.05.

### Identifying AD biomarkers and drug targets from the literature

A systematic review of the literature was conducted in PubMed on February 10, 2026. The search strategy included the following terms: plasma, cerebrospinal fluid, cortex, brain, and proteomics and were limited to titles and abstracts. Duplicate pre-print and published version PMIDs were removed. The remaining tiles, abstracts, and full texts were rated based on *a priori* eligibility criteria (Table 3).

**Table 3.** Selection criteria for studies included in the systematic literature review.

|  |
| --- |
| **Inclusion criteria** |
| Participants with Alzheimer's disease |
| Used high-throughput proteomics |
| Human plasma, CSF, or cortex samples |
| Listed specific protein biomarkers and/or drug targets |
| Published pre-print or manuscript |
| Primary study |
| Written in English |

Studies were excluded on the basis of (1) participants with diseases other than AD or limited to specific genotypes, (2) did not measure proteins using a high throughput method (e.g. mass spectrometry), (3) no analyses of human samples, (4) not identifying specific protein biomarkers and/or drug targets (e.g. modules rather than proteins), (5) describing method developments rather than primary results, and (6) not in English. Reviews were identified during the screening of abstracts and were removed prior to assessing full texts. For each included study, we extracted the proteins that the authors highlighted as candidate AD biomarkers or drug targets. A full list of included studies is included in the Supplementary References and PMIDs in Supplementary Table 2.

---

**Supplementary Figure 1.**
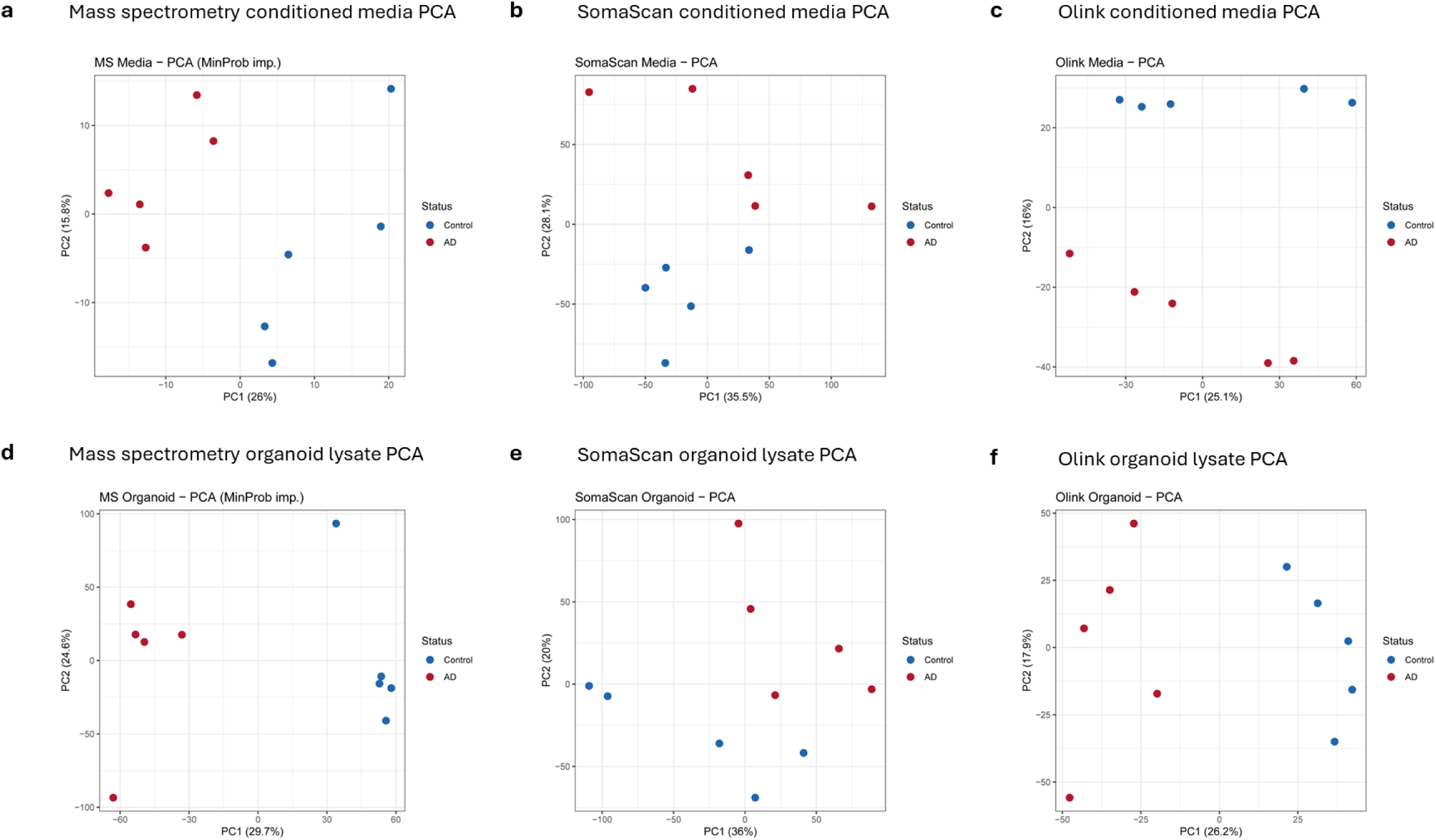
Differentially abundant proteins separate AD and control organoids across all platforms and sample types. Principal component analysis (PCA) of DAPs between AD and control organoids for each of the six platform and sample type combinations, showing separation by disease status in every case. (**a-c**) Conditioned media profiled by (**a**) LC/MS, (**b**) SomaScan, and (**c**) Olink, respectively. (**d-f**) Organoid lysate profiled by (**d**) LC/MS, (**e**) SomaScan, and (**f**) Olink, respectively. Abbreviations: AD: Alzheimer’s disease; DAP: differentially abundant protein; LC/MS: liquid chromatography mass spectrometry; PCA: principal component analysis.

**Supplementary Figure 2.**
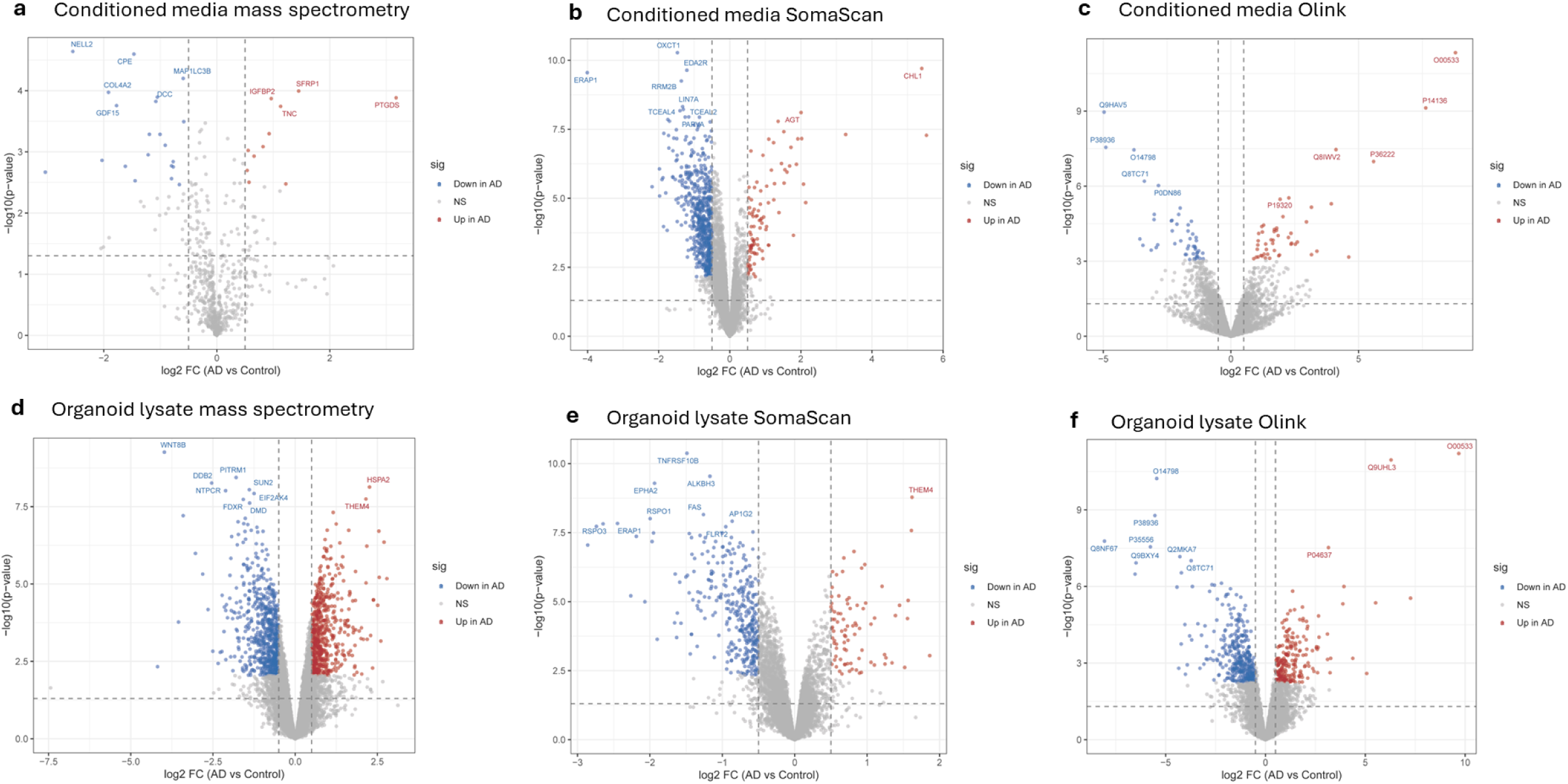
Differentially abundant proteins in the organoid model across all platforms and sample types. Volcano plots of conditioned media DAPs across (**a**) LC/MS, (**b**) SomaScan, and (**c**) Olink. Volcano plots of organoid lysate DAPs across (**d**) LC/MS, (**e**) SomaScan, and (**f**) Olink.

**Supplementary Figure 3.**
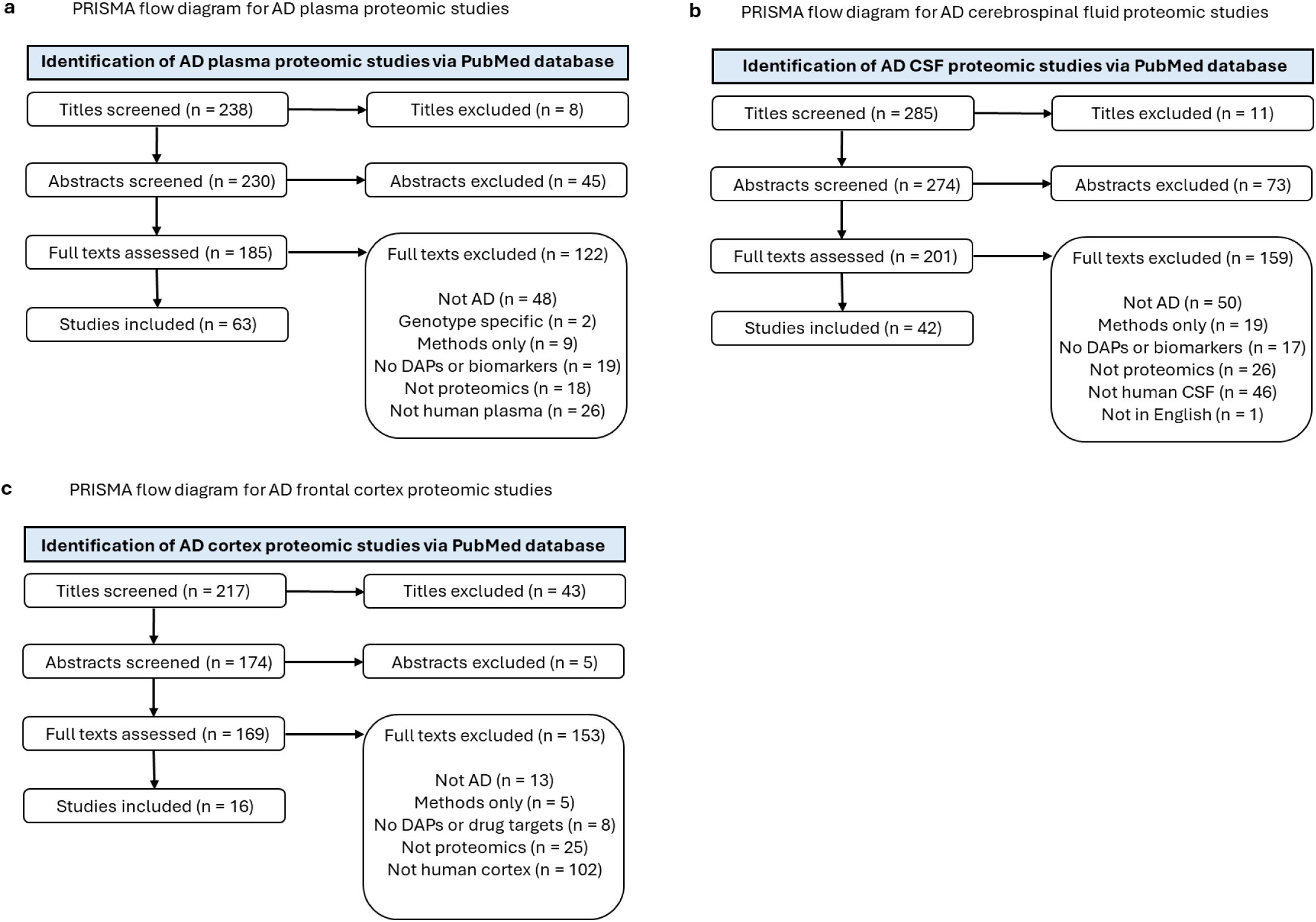
Systematic literature review of studies identified AD biomarkers and drug targets from clinical proteomics studies. PRIMA flow diagrams of the selection process for (**a**) plasma, (**b**) CSF, and (**c**) frontal cortex clinical proteomics studies. Abbreviations: Abbreviations: AD: Alzheimer’s disease; CSF: cerebrospinal fluid; DAP: differentially abundant protein.

**Supplementary Figure 4.**
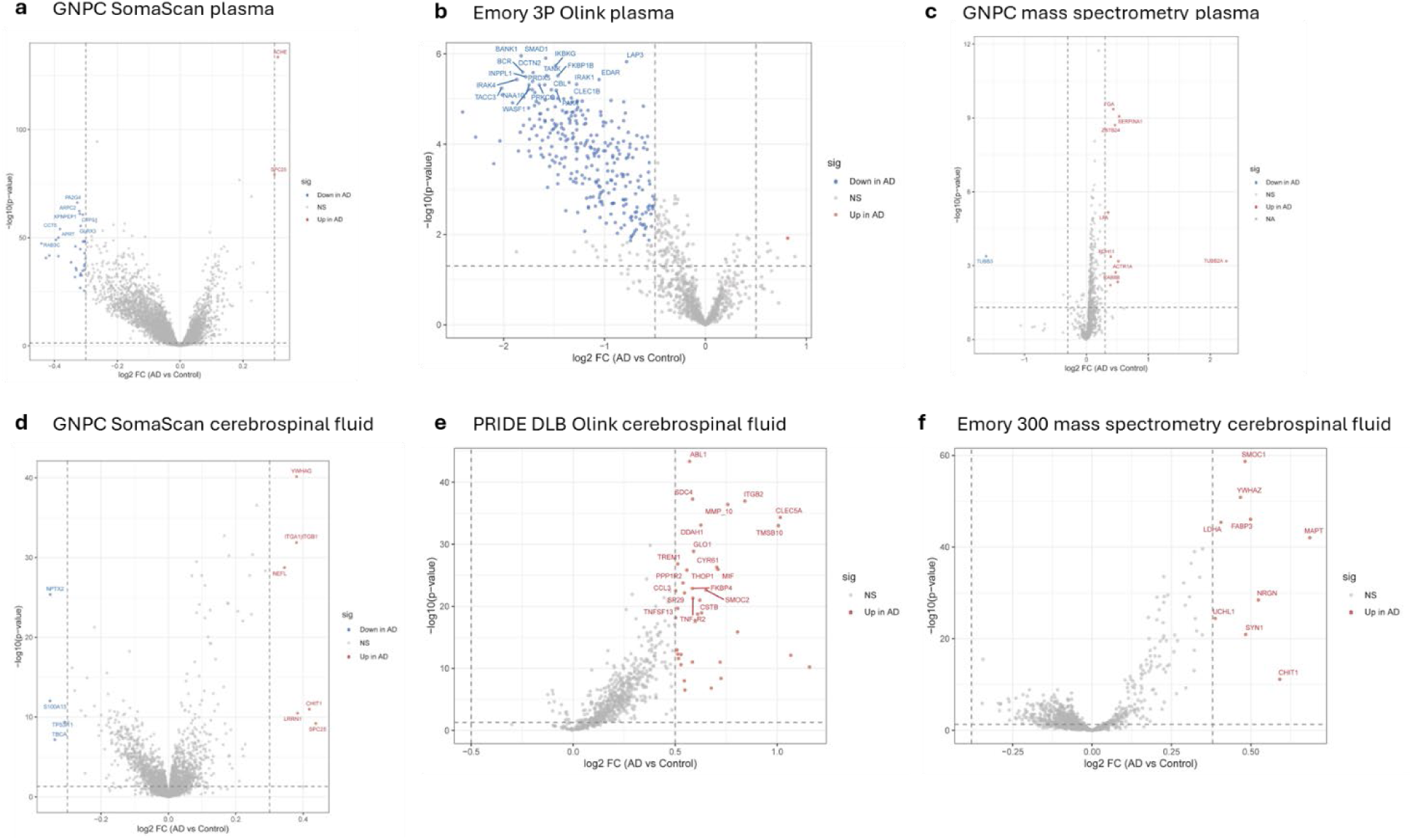
Differentially abundant proteins in the clinical cohorts across plasma and CSF on all platforms. Volcano plots of plasma DAPs across (**a**) SomaScan, (**b**) Olink, and (**c**) LC/MS. Volcano plots of CSF DAPs across (**d**) SomaScan, (**e**) Olink, and (**f**) LC/MS.

