## Supplementary References for "Patient cerebral organoids capture Alzheimer’s disease proteomic biomarkers and drug targets"

Plasma proteomic studies included in the systematic literature review ^1-63^

Cerebrospinal fluid proteomic studies included in the systematic literature review ^4,11,40,64-102^

Cortex proteomic studies included in the systematic literature review ^11,103-117^
